# Open source 3D printed Ventilation Device

**DOI:** 10.1101/2020.05.21.108043

**Authors:** Ahmad Faryami, Carolyn A Harris

## Abstract

COVID-19 is an acute respiratory tract infection caused by a coronavirus known as SARS-CoV-2. The common signs of infection include respiratory symptoms such as shortness of breath, breathing difficulties, dry cough, fever, and in some patients, severe acute respiratory syndrome, kidney failure, and death. 312,009 deaths from COVID-19 has been reported as of today. While respiratory symptoms are commonly caused by the infection, the use of mechanical ventilation is required for some patients. The following is intended to review the development and testing of a 3D printed and open-source mechanical ventilation device that is capable of adjusting breathing rate, volume, and pressure simultaneously and was designed according to the latest clinical observations of the current pandemic. The intuitive design of this device along with the use of primarily 3D printed or readily available components allow the rapid manufacturing and transportation of this ventilation device to the impacted regions.

## Background

According to the World Health Organization, more than 4,628,903 cases of COVID-19 infections have been confirmed around the world^1^. However, many experts estimate the actual number of patients to be significantly higher than the reported cases. Although many aspects of the infection are currently unknown, health organizations, pharmaceutical companies, and researchers are rapidly studying the infection^2^. The preliminary clinical data collected in China by Dr. Wei-jie Guan and his colleagues indicate that 1.4% of the admitted patients died from the infection. However, 41.3 % of the patients received oxygen therapy and mechanical ventilation was administered to 6.1% of patients^3^.

These data are consistent with the clinical observations from Iran and Italy. While the number of confirmed cases of the infection has been exponentially rising around the world, the shortage of medical supplies and equipment is one of the main challenges in providing effective treatment to the patients^4–6^.

Open-source medical technologies have been developed for testing or experimental use in the past, but the global pandemic has elevated the importance of open-source technologies and medical equipment to rapidly expand the capacity of the healthcare system^7^.

This open-source positive pressure ventilation device (OSPPVD) is a response to the shortages in the hospitals’ ventilation capacity, which has been reported to be essential to many COVID-19 patients throughout the world. OSPPVD has been designed according to health care professionals’ descriptions and observations of COVID-19 infection instead of recreating already existing technologies^5,8,9^. However, it is not intended to replace nor negate the importance of medications as well as care provided by healthcare professionals. The open-source ventilation device is intended to assist in increasing the capacity of healthcare centers or provide the healthcare providers the tools necessary to provide effective care to patients^10^.

## Methods and Materials

This device operates based on the changes in hydrostatic pressure within three partially empty and interconnected containers. This is achieved through lowering and raising one of the containers relative to the other two stationery containers. At equilibrium, the height of the water column is equal in all three containers (Figure 1). The change in the height of the actuating container results in a change in hydrostatic pressure and displacement of air and water (Figure 2-3).

**Fig. 1.**
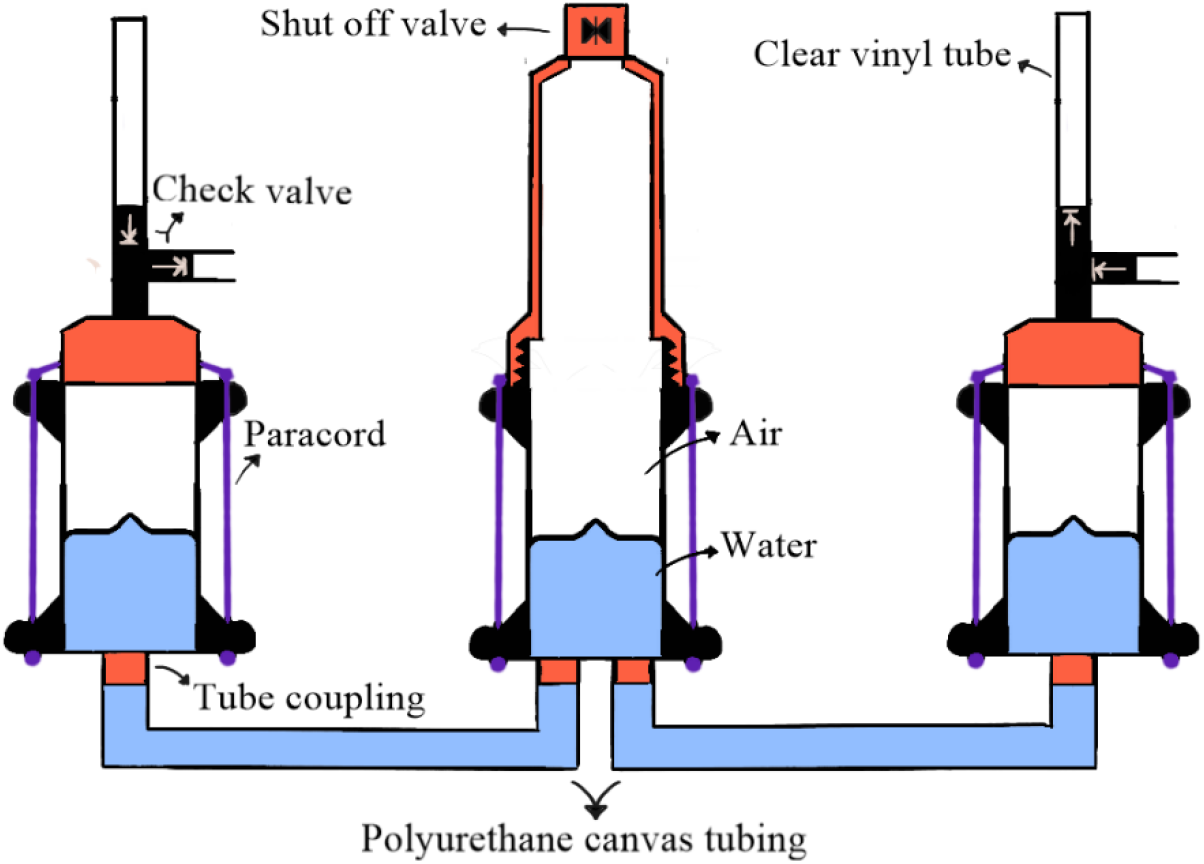
The starting position of the three interconnected containers relative to each other is shown in this figure. At this stage, water is level in all three containers. The two stationary containers are identical in capacity and construction except for the direction of the check valves indicated above.

**Fig. 2.**
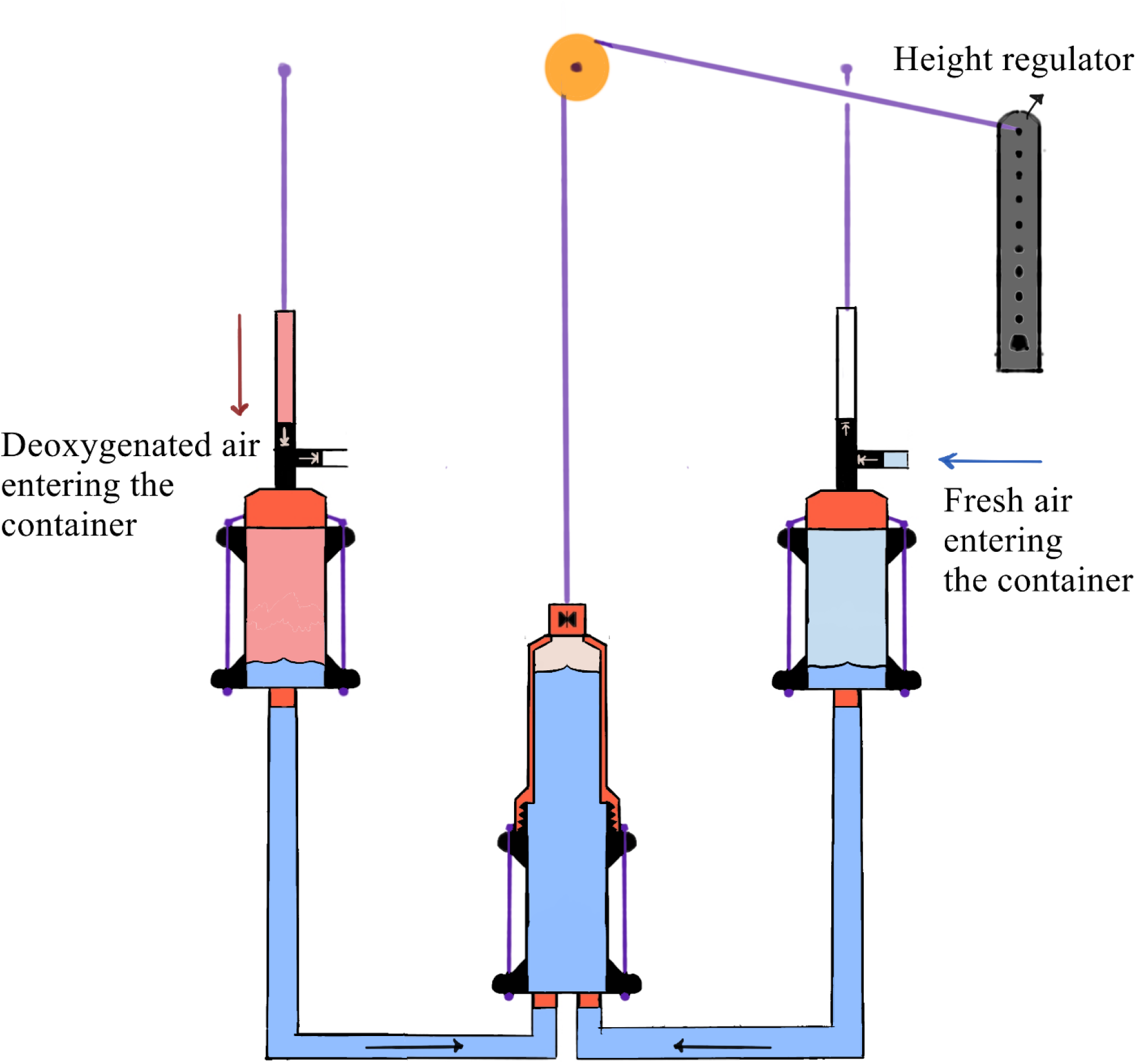
The simultaneous intake of fresh oxygenated air from a secondary source, oxygen tank, or atmosphere, and the intake of deoxygenated air from the lungs is shown in this figure. The lowering of the actuating container relative to the stationary containers result in an inflow of water and displacement of air.

**Fig. 3.**
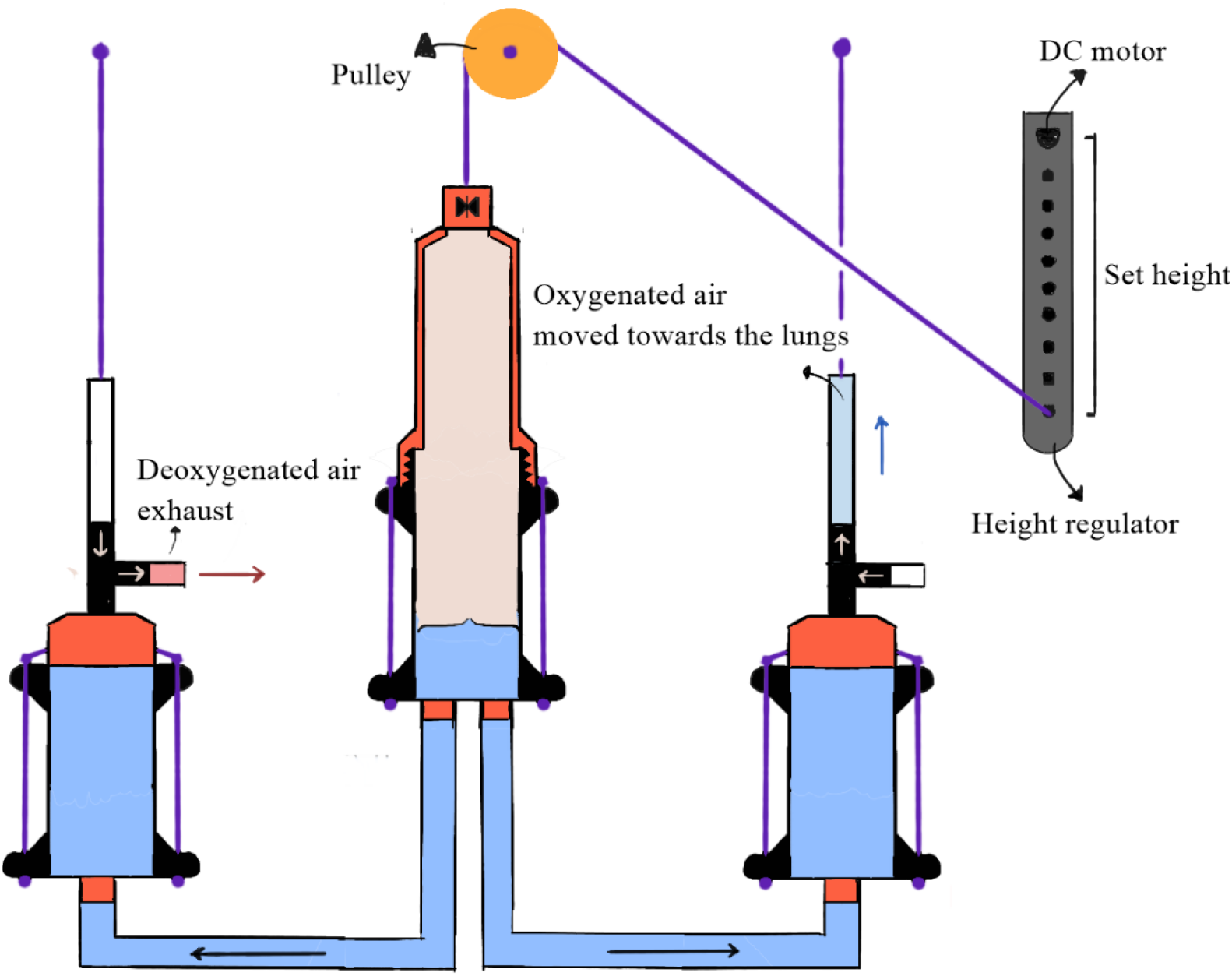
The containers are shown during the air compression half-cycle. The flow of water from the actuating container to the stationery containers compresses the entrapped air within the containers. The fresh oxygenated air is moved towards the lungs while deoxygenated air is expelled out of the container

The flow of water from one container to the other two containers results in the displacement of air transferred to the patient through a series of tubes and one-way check valves. In this setup, two of the containers were secured at a set height while the other container was attached to an electric motor and was lowered or raised relative to the other containers. This setup is particularly useful since it does not require traditional compression equipment such as pistons or precision parts such as gearboxes while a low number of moving parts would allow relative ease in sterilizing and reusing the equipment between patients.

### Electric Motor

A geared DC motor (Amazon, Seattle, WA, USA) rated at 30 RPM at a 12V input was utilized in this setup while the frequency of rotation translated to one cycle of inhalation and exhalation. The input voltage to the motor was adjusted via a DC motor speed controller (Onyehn, Seattle, WA, USA) in order to achieve the required respiration rate. A Neiko 20713A laser tachometer (Amazon, Seattle, WA, USA) was used to measure and record the revolution rate. A 3D printed motor housing was designed to retain the motor on a tripod stand mounted on the wall using two screws as illustrated in figure 4.

**Fig. 4.**
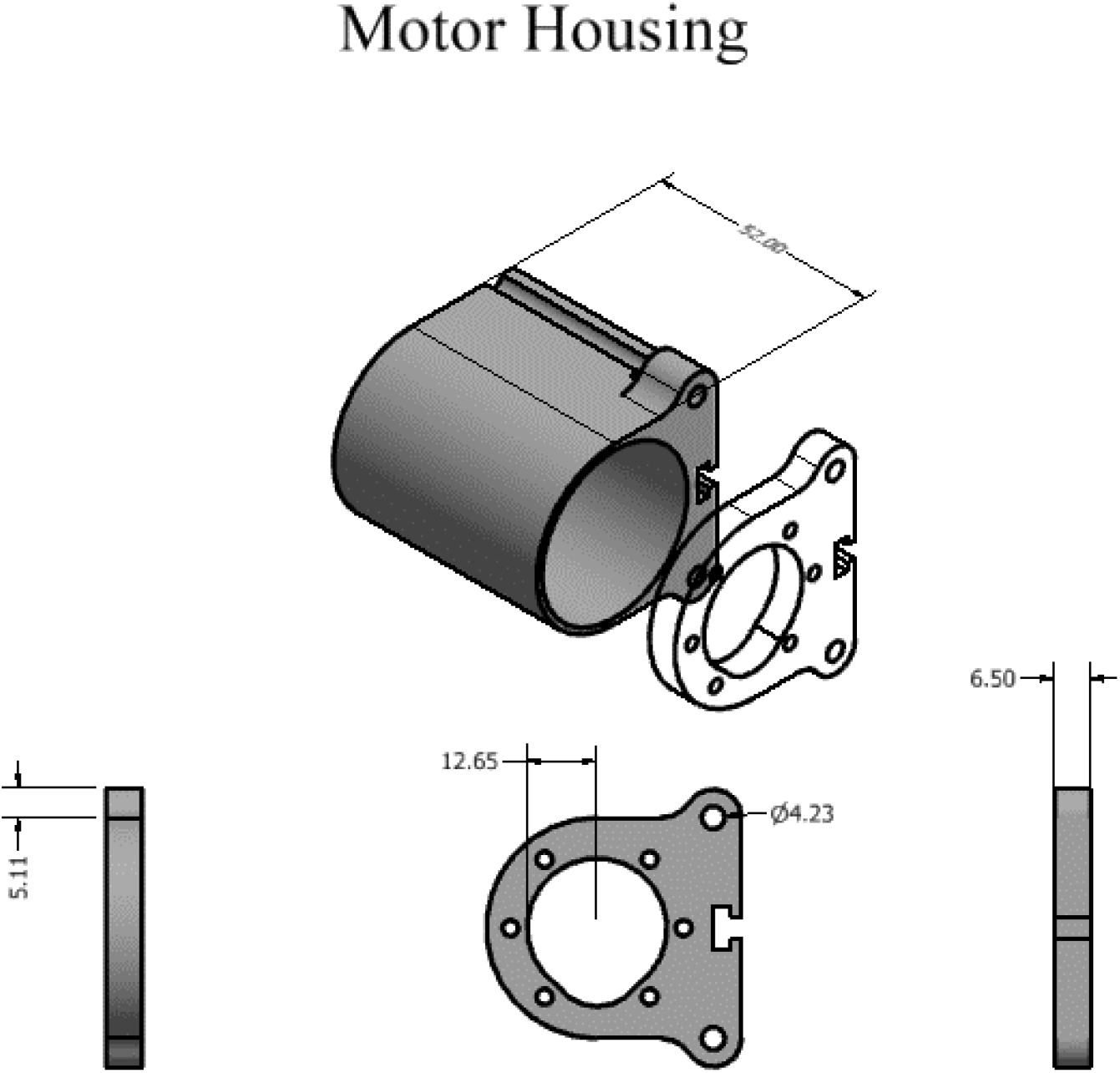
Wall-mounted motor housing made of two separate pieces which are designed to secure the motor once mounted on a vertical or horizontal surface

**Fig 5.**
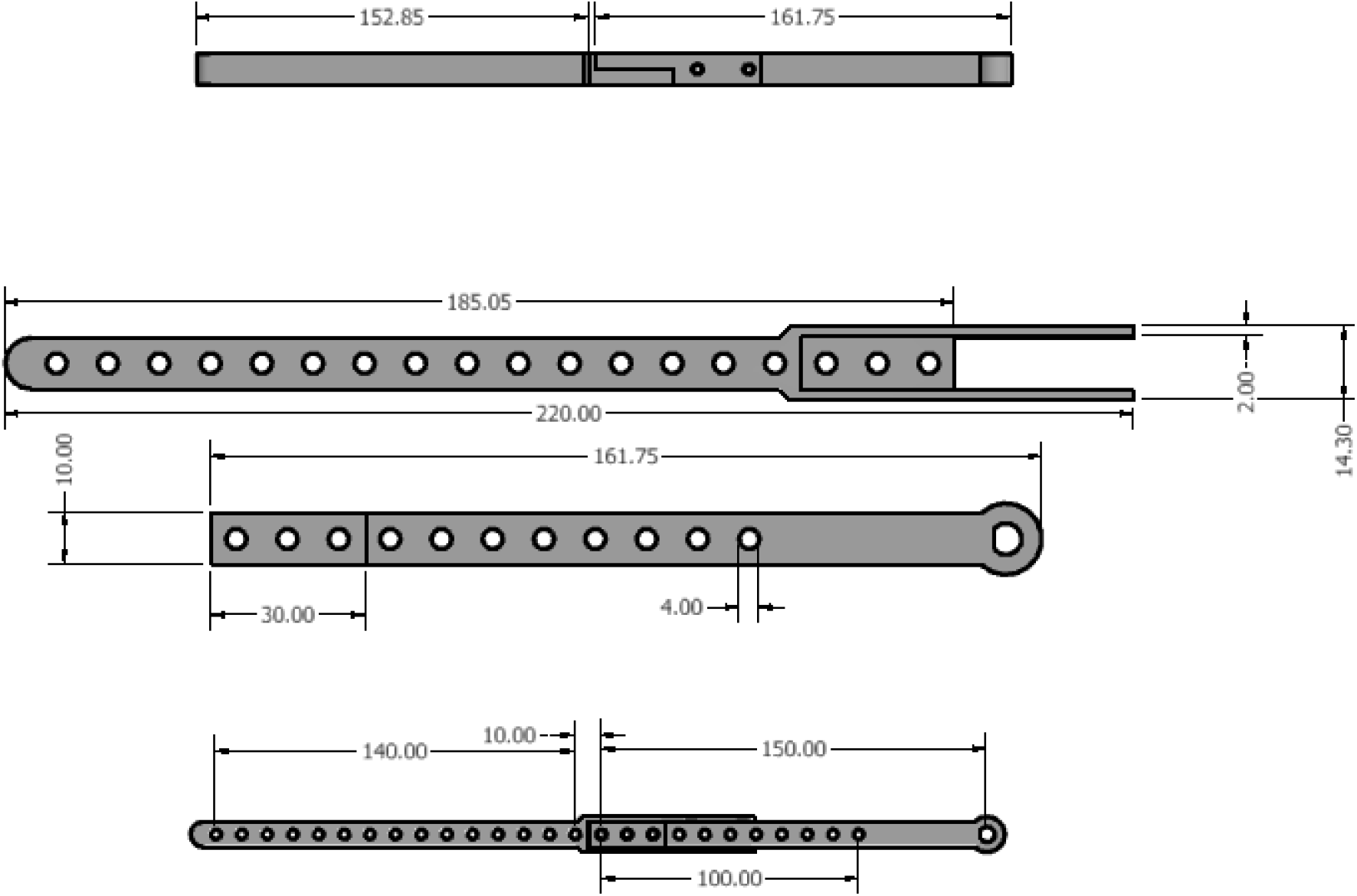
Height regulator with adjustable length ranging from 50 to 3000 mm.

### Height Regulator

Importantly, the height regulator translates the rotational output of the electric motor, then subsequently to linear motion to raise and lower the container. Since the relative height of the containers is the variable factor of hydrostatic pressure in this setup, controlling the height difference between the containers determines the set inhalation and exhalation pressure. Tidal volume is determined by the volume of water displaced during each cycle as a function of the height difference. The height regulator is 3D printed in two parts to make sure most printers with a building surface of at least 22 cm in any dimension could also manufacture the parts. Finally, a shut off valve was incorporated in the device to adjust tidal volume and to avoid overdistention of normal alveoli and volume-induced trauma.

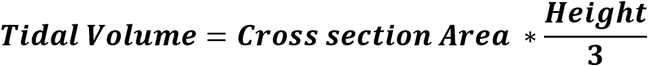

**Equitation 1. Calculating the maximum tidal volume using the height set by the regulator and the cross-sectional area**

### Pulley System

Although the inherent advantages of a rigid metal frame with linear rails were initially put into consideration for this device, a 3D printed pulley system was eventually constructed instead, to reduce the overall number of all the required parts, cost and to improve the ease of storage and transport. A 3D printed pulley system was constructed using paracord, a 22 mm skateboard ball bearing, and 3D printed parts. Around three meters of readily available paracord was used in this setup.

### Valves and Tubing

Different types of tubing were utilized in the construction of this project (Table 1). The tubing is used to transfer water or air. Four 13mm OD check valves were used in this setup to carry oxygenated air to the patient and carry the expired air away. The tubing that was used to interconnect the containers has a wider inner diameter to allow equilibrium to be reached rapidly between the containers.

**Table 1.** List of the tubing used in the device and the total length

| Type of Tubing | Inner Diameter<br>(mm) | Outer Diameter<br>(mm) | Length (mm) |
| --- | --- | --- | --- |
| Polyurethane Canvas | 25.4 | 27.5 | 500 |
| Clear Vinyl | 13 | 16 | 300 |

### 3D Printing

All parts were printed on an Anycubic Mega (Amazon, Seattle, WA, USA) using OVERTURE 1.75mm Polylactic acid filament (Amazon, Seattle, WA, USA). A 0.6 mm nozzle was used for printing the files instead of 0.4 mm nozzle to reduce the total printing time (Table 3). All the Computer-Aided Design files (CAD) were made using Autodesk Inventor 2020 software and all the parts were printed without any additional supports, or brim. The 3D printed parts (Figure 6) were color-coordinated with schematic diagrams (Figure 1-3).

**Fig. 6.**
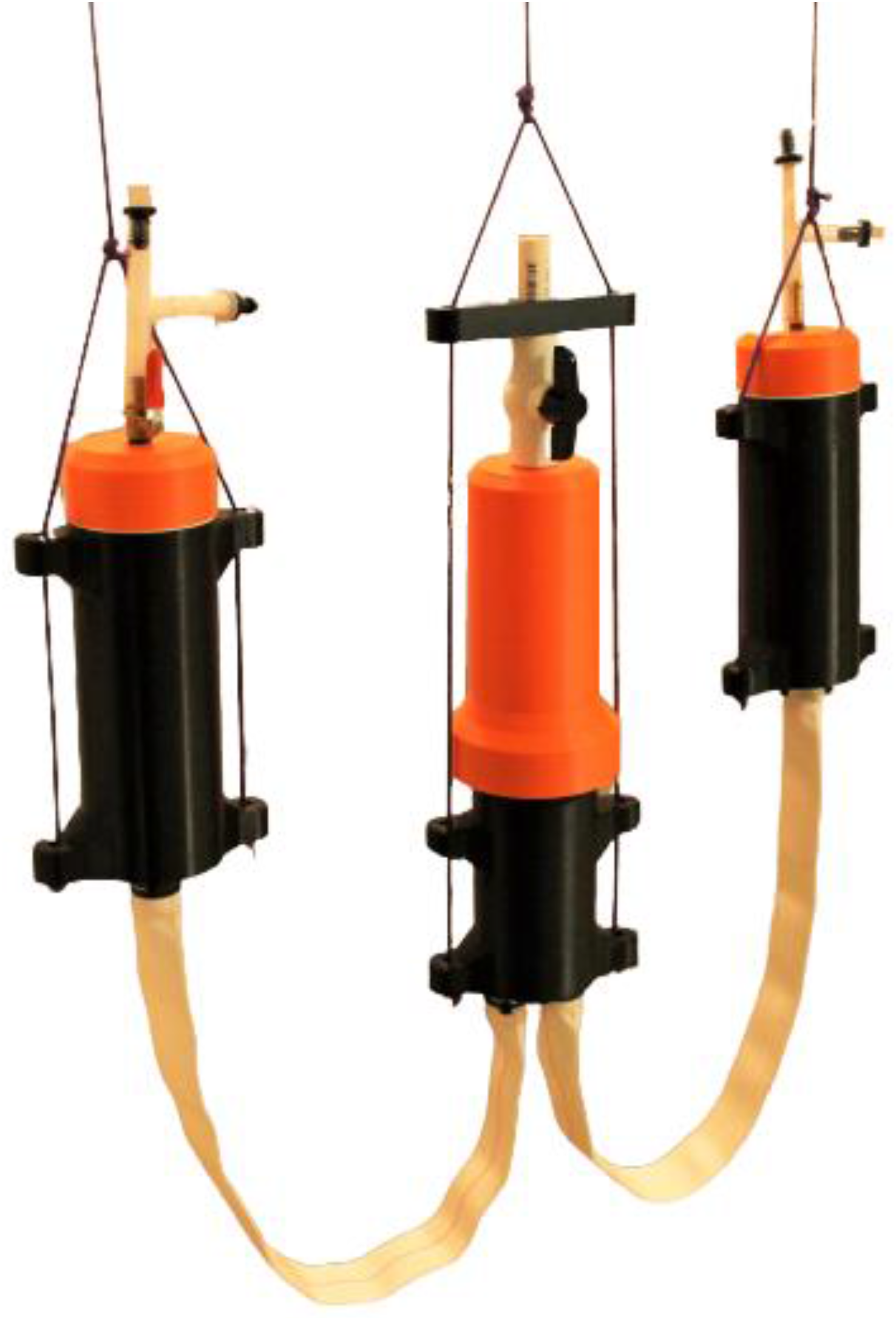
3D printed ventilator assembled prior to testing pressure and ventilation rate

**Fig. 7.**
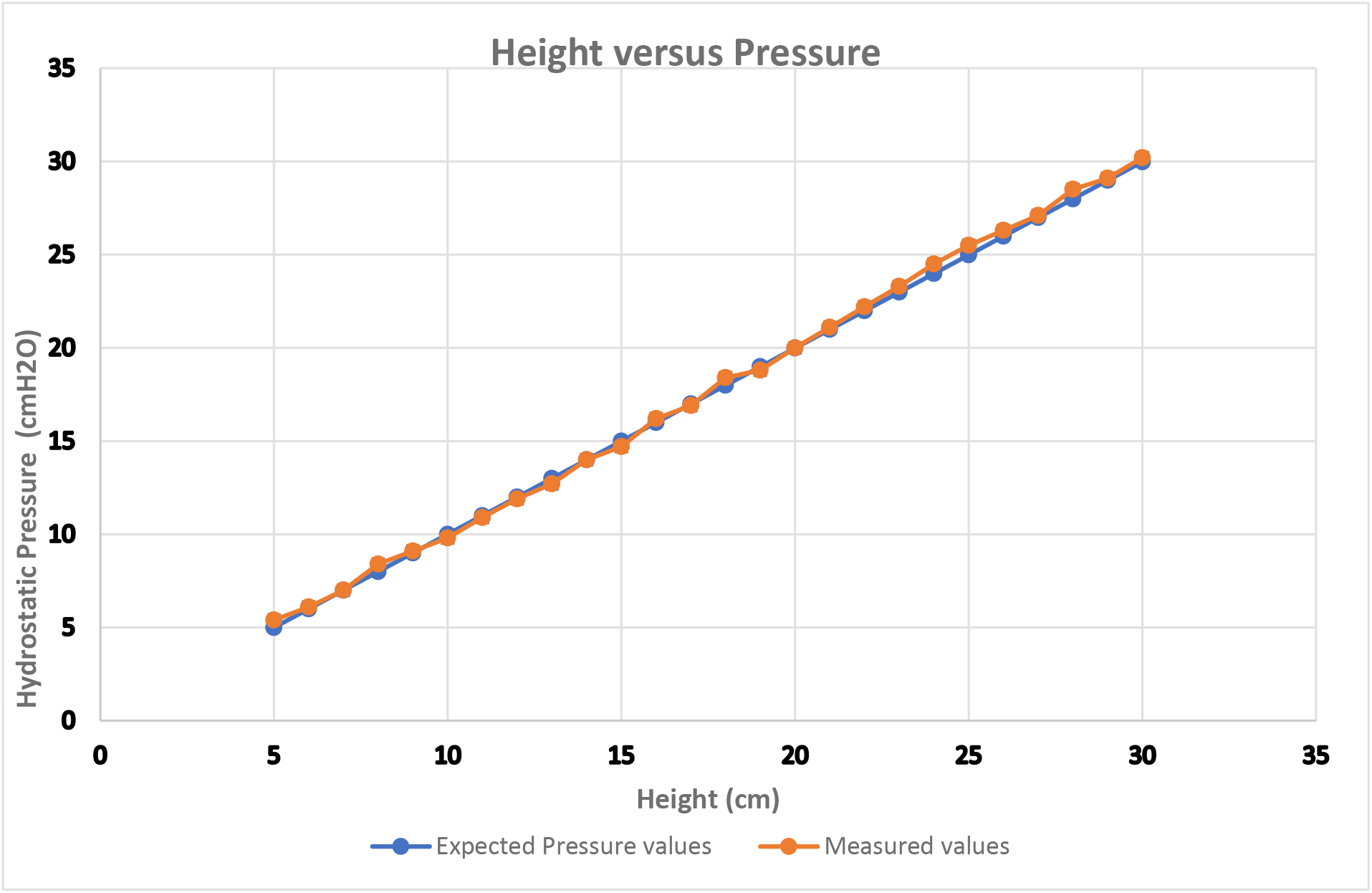
Illustrates the relationship between pressure and height while demonstrating expected and measured values

### Containers

The 3D printed containers were specifically designed to be watertight and the internal walls were optimized for fused deposition manufacturing process to avoid water or air leakage. The inlet and outlet joints were designed at a reasonable tolerance to be printable with almost all consumer-grade 3D printers while eliminating the need for adhesives in order to further minimize the chance of leakage while in operation and reduce assembly time.

### Statistical Analysis

In this experiment, the OSPPVD was operated over a 24-hour period to evaluate the device performance as well as measuring the output pressure. All the pressure measurements were made using the Hti-Xintai dual-port handheld digital manometer (Amazon, Seattle, WA, USA). All pressure measurements were recorded by a digital camera at 30 frames per second and the pressure data from the recording were extracted through processing the video at a frame by frame basis using Free Video to JPG Converter software.

## Results

The systematic output is a direct result of precision and accuracy of the achieved height by the actuating container. While the use of a pulley system instead of a solid frame was determined as a cost reductive measure and significantly increased the mobility of the setup, the lack of a rigid framework also increased the difference between the set and measured height (Table 2).

**Table 2.** Summary of actuator travel distance as well as measured deviation from the expected values

| **Range of travel distance** | **Maximum measured travel distance** | **Minimum distance travelled** | **Maximum deviation from expected value** | **Minimum deviation from expected value** |
| --- | --- | --- | --- | --- |
| **5-30 cm** | 30.3 cm | 4.9 cm | $\pm 0.4$ cm | 0 cm |

Despite the micro-gaps that occur during the additive manufacturing process, no considerable leakage was observed during the experiment. Though other polymers such as High Impact Polystyrene (HIPS) or Polyethylene terephthalate (PETG), are better suited for this application, the ease of use and ubiquity of Polylactic Acid filament allowed rapid prototyping and iterating of this device. Optimization of printing speed and quality show exact printing settings vary between different makes and models (Table 3). While the steady increase in pressure observed by a gradual pressurization is best represented by a sinusoidal wave, the handheld digital manometer was not capable of providing a sufficient passage for the flow of large volume air displaced to measure pressure changes through a cycle. However, the pressure achieved at each height was measured and the expected pressure was calculated using the hydrostatic pressure formula. Regular tap water at room temperature was used in this experiment.

**Table 3.** 3D printing results.

|  |  |
| --- | --- |
| **Total 3D printed weight** | 1275 grams |
| **Material** | Polylactic acid (PLA) |
| **Bilateral tolerance** | 0.2 mm |
| **Assembly time** | 1 hour and 45 minutes |
| **Total printing time** | Approximately 42 hours |
| **Layer height** | 0.2 mm |
| **Printing speed** | 75 mm/sec |
| **Nozzle diameter** | 0.6 mm |

While a maximum of ±0.5 *cmH*_2_*O* variance between the height set by the height regulator and the pressure measured by the digital manometer was observed during this experiment, the height regulator may also be used as an indicator for the peak pressure in *cmH*_2_*O* assuming that the density of the medium is known.

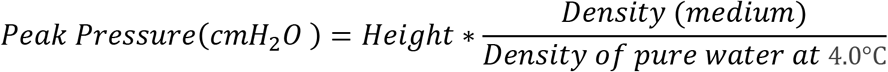

**Equation 2. Approximating peak pressure using the height set by the height regulator**

## Discussion

The slight difference between the measured and expected pressure is within the range of difference observed in expected and measured traveled distance. While paracord is a readily available and cost-effective material for building the pulley system, especially due to its high tensile strength. However, stretching in length under load was observed which results in a deviation from the expected height and pressure. A sturdier material may be more suitable for this application.

Clinical observations indicate that maintaining a positive end-expiratory pressure (PEEP) is crucial in avoiding alveolar collapse among COVID-19 patients. Maintaining PEEP is easily achieved through a secondary source of compressed air and a pressure-regulated shut-off valve. However, this setup requires further studies and is currently under development^9,11^.

The steady increase and decrease in pressure are best fit to a sinusoidal graph that is similar to the breathing pattern observed in human adults. The gradual decrease in pressure could be beneficial in avoiding pressure related injury that may result in scar formation and fibrosis. According to the initial reports published by healthcare providers, the loss of compliance due to the onset of fibrosis in the lungs is observed in patients diagnosed with severe cases of COVID-19 infection^9^. An advantage of OSPPVD presented here is its capability to perform hyperventilation with high flow rates under relatively wide pressure range from 5 *cmH*_2_*O* that would allow the delivery of oxygen without the over-expansion of the lungs while being able to supply pressurized air as high as 40 or 50 *cmH*_2_*O* even though the current setup is designed to only reach 30 *cmH*_2_*O*.

The underlying functioning principle of this setup allows it to run through many cycles of inhalation and exhalation with minimal cyclical system fatigue (wear and tear). To our knowledge of the currently available systems, this is an advantage over other open-source ventilators that rely on the compression of a flexible air chamber with limited compression cycles before it has to be replaced from loss of compliance. Furthermore, the flexible air chamber is best used when providing positive pressure necessary during the inhalation. This method of ventilation might not be suitable for patients with COVID-19 as they mainly rely on lungs’ compliance for exhalation. Fibrosis and the loss of lung compliance is a common symptom among COVID-19 patients and may not allow the patient to exhale normally^12^.

A conscious effort was made to reduce the number of electronic components needed to manufacture and control the device to reduce the production cost as well as minimizing the chance of device malfunction and injury to the patient due to electronic or software malfunction. However, another iteration of this setup is currently under development that is controlled by a micro-controllers and sensors to adjust the breathing rate and the intake volume according to the patients’ needs autonomously and in real-time.

Oxygen therapy is an essential treatment for many infected patients, and this setup was designed to be capable of intaking concentrated oxygen and supplying the patient according to the healthcare provider’s specified volume and pressure. This may be achieved through a regulated cylinder and an oxygen tank connected to the intake port of the positive pressure container. Moisturizing and warming up the intake air are another feature that was put into consideration while designing this device. During this experiment, the intake tube was extended below the waterline in the compression chamber which allowed the air to percolate through warm water before being compressed out of the container. The container was 3D printed from Polyethylene terephthalate (PETG), due to its thermal properties compared to PLA^13^. This method was used to make the intake air warm and humid before being compressed and expelled. Although an increase in the air temperature was noticed during this experiment, further analysis is required to measure the moisture content and confirm these results. Furthermore, the exhaust air from the negative pressure container may contain airborne water droplets that could be hazardous to the medical professionals as well as other people in the vicinity of the patient. The traditional methods of spreading airborne viruses such as fine filters may be used in this setup. A similar method may also be utilized to remove contaminants before it is transported to the lungs^14^.

Although calculating the tidal volume using the provided equation is rather simple, it is still necessary to include an inline pressure-regulated valves in order to avoid any injuries due to over-expansion of the lung. Other safety valves and sensors may be added in order to protect the patient against ventilator-induced injury.

## Funding

Funding was provided by the College of Engineering, Department of Chemical Engineering and Materials Science at Wayne State University.

## Notes

### Competing Interest Statement

The authors have declared no competing interest.

## References

1. Organization, W. H. Coronavirus disease 2019. https://www.who.int/emergencies/diseases/novel-coronavirus-2019.

2. Wölfel, R. et al. Virological assessment of hospitalized patients with COVID-2019. Nature (2020) doi:10.1038/s41586-020-2196-x.

3. Guan, W. J. et al. Clinical Characteristics of Coronavirus Disease 2019 in China. N. Engl. J. Med. (2020) doi:10.1056/NEJMoa2002032.

4. Khosrawipour, V. et al. Failure in initial stage containment of global COVID‐19 epicenters. J. Med. Virol. 1–5 (2020) doi:10.1002/jmv.25883.

5. Rokni, M., Ghasemi, V. & Tavakoli, Z. Immune responses and pathogenesis of SARS-CoV-2 during an outbreak in Iran: Comparison with SARS and MERS. Rev. Med. Virol. (2020) doi:10.1002/rmv.2107.

6. Carenzo, L. et al. Hospital surge capacity in a tertiary emergency referral centre during the COVID-19 outbreak in Italy. Anaesthesia 1174, 1–7 (2020).

7. Pearce, J. M. A review of open source ventilators for COVID-19 and future pandemics. F1000Research 9, 218 (2020).

8. Sorbello, M. et al. The Italian coronavirus disease 2019 outbreak: recommendations from clinical practice. Anaesthesia 724–732 (2020) doi:10.1111/anae.15049.

9. Iwasawa, T. et al. Correction to: Ultra-high-resolution computed tomography can demonstrate alveolar collapse in novel coronavirus (COVID-19) pneumonia (Japanese Journal of Radiology, (2020), 10.1007/s11604-020-00956-y). Jpn. J. Radiol. 38, 399 (2020).

10. Clarke, A. L., Stephens, A. F., Liao, S., Byrne, T. J. & Gregory, S. D. Coping with COVID ‐19: ventilator splitting with differential driving pressures using standard hospital equipment. Anaesthesia 1–9 (2020) doi:10.1111/anae.15078.

11. Singh, A. G. & Chaturvedi, P. Clinical trials during <scp>COVID</scp> ‐19. Head Neck hed.26223 (2020) doi:10.1002/hed.26223.

12. Sun, P., Lu, X., Xu, C., Sun, W. & Pan, B. Understanding of COVID-19 based on current evidence. J. Med. Virol. 92, 548–551 (2020).

13. Szykiedans, K., Credo, W. & Osiński, D. Selected Mechanical Properties of PETG 3-D Prints. Procedia Eng. 177, 455–461 (2017).

14. Liu, D. C. Y. et al. Adapting reusable elastomeric respirators to utilise anaesthesia circuit filters using a 3D‐printed adaptor; a potential alternative to address N95 shortages during the COVID‐19 pandemic. Anaesthesia anae.15108 (2020) doi:10.1111/anae.15108.

